# From Encoding To Conscious Report: Electrophysiological Signatures Of Iconic Memory Revealed By A Partial Report Task

**DOI:** 10.64898/2026.03.13.711298

**Authors:** D. Bonfanti, S. Mele, E. Bertacco, C. Mazzi, S. Savazzi

## Abstract

Despite numerous investigations, a comprehensive electrophysiological characterization of iconic memory remains lacking. Through a partial report paradigm, we aimed to shed light on this topic by disentangling electrophysiological activity related to stimulus perception from that linked with the specific task. We collected EEG data from 26 participants while they performed a partial report task. They were shown circular arrays of six letters lasting 100 ms. After the stimulus, an acoustic cue instructed the participant to report on which side of the array. Differences between reporting conditions were primarily evident in the time window 850-1100 ms, characterized by a positive component predominantly over parieto-occipital electrodes ipsilateral to the reporting side. Through linear regression, we also found a positive relationship between P1 and participants’ accuracy, as well as negative relationships between P3, VCR, TIF, and accuracy. Our results provide an overview of the different processes involved in iconic memory, corroborating the distinction between a series of neural mechanisms responsible for encoding and maintaining the entire stimulus and higher-order processes in charge of selecting an information subset for conscious report. The TIF component, in particular, could act as a key filtering mechanism to prevent irrelevant information from being selected for further processing. Our results provide, for the first time, a thorough characterization of the electrophysiological dynamics behind iconic memory. Moreover, implications for the consciousness debate are discussed, particularly regarding the overflow argument and how our results could be read through its lens.

## Introduction

Iconic memory (Neisser, 1967) is a kind of visual, short-term memory store in which perceptual information persists after the disappearance of the physical stimulus. Sperling, in his seminal work, was the first to identify and study the properties of this memory storage extensively (Sperling, 1960). In a series of experiments, he presented participants with arrays of letters and numbers of varying sizes for different durations. Participants then had to make a report in one of two possible modalities: either give a full report, trying to correctly indicate all the symbols displayed, or a partial report (Chow, 1985), specifying the symbols of only one row, cued immediately after stimulus offset by an acoustic tone. Results from full reports were in accordance with a classical working memory interpretation: participants could report an average of four letters, a number that remained constant even at longer stimulus duration, and that was consistent with the idea of an upper limit in working memory capacity. Partial report results, on the other hand, did not fit in this framework, providing evidence for the existence of another storage mechanism preceding working memory. When asked to report only a single row, indeed, participants were able to indicate all the relevant letters even with stimuli lasting 50 ms: this suggested that participants had access to the visual representation of the whole stimulus, from which they could extract and report with great accuracy the cued subset even when it was not visible anymore. Sperling also found that the content of this early-stage, high-capacity sensory buffer – corresponding to nowadays iconic memory – quickly decayed over time, with just a few milliseconds of delay between stimulus offset and cue onset being enough to decrease participants’ performance to the full report condition.

Over the years, the advantage of iconic memory in partial reports has been replicated several times, although to different extents (Bradley-Garcia et al., 2023). Numerous experiments have investigated the functioning and characteristics of iconic memory, trying to elucidate its mechanisms. In particular, the relationship with attention (Bachmann & Aru, 2015; Mack et al., 2015, 2016; Persuh et al., 2012), the impact of training (Gong et al., 2024), the degree of integration with subsequent visual percepts (Sugita et al., 2018), the nature and characteristics of the stimulus (Di Lollo, 1984; Graziano & Sigman, 2008; Haun & Tononi, 2025; Sakitt & Appelman, 1978), the nature of decay (Pratte, 2018), the modulation elicited by neurodevelopmental and neuropathological conditions (Gosavi & Hubbard, 2019; Hahn et al., 2011; Lu et al., 2005) are just some of the different aspects of iconic memory that have been the focus of experimental research.

Despite this extensive investigation, efforts to ascertain the electrophysiological and neural correlates underlying this sensory storage have been less consistent. From a source perspective, electrophysiological studies conducted on macaque monkeys and fMRI studies conducted on humans have identified potential neural correlates either in the primary visual cortex (Ruff et al., 2007; Teeuwen et al., 2021) or in higher inferotemporal visual areas (Keysers et al., 2005), with the exact location possibly influenced by the degree of complexity of the presented stimulus. However, a precise characterization of the electrophysiological profile of iconic memory is still lacking. It is known that the elaboration of visual information begins with the processing of elementary visual features, which are assembled to internally reconstruct the original image as an icon, which outlasts the external stimulus; this icon can then be employed by subsequent cognitive operations. While many different operations are directly involved in the build-up of an icon and its subsequent manipulation, from an electrophysiological perspective there is no clear separation between the processes associated with stimulus perception and its storage in iconic memory, and those related to the post-perceptual processes necessary to create a response output. To investigate this distinction, the ideal task would consist of identical visual stimuli followed by different task demands, thereby allowing the isolation of iconic memory activity from subsequent processing. In this respect, partial report is well-suited for this experimental situation: while the characteristics of the presented stimulus are consistent across conditions, this paradigm allows modulating the information subset to be reported. This dissociation provides for modulating post-perceptual processes while keeping perceptual processes constant, shedding light on the actual iconic memory mechanisms and separating them from response mechanisms.

In the present study, therefore, we adopted a partial report paradigm while recording the EEG signal. Participants saw a circular array of letters, and an auditory cue presented after stimulus disappearance instructed them to report either the left or the right half of the array. This allowed us to record activity associated with perception and maintenance of the perceptual image, and to dissociate it from that related to response preparation and post-perceptual processing.

## Materials and Methods

### Participants

A total of 26 volunteers (15 F, mean age 25.6 ± 4.7 years, right-handedness ascertained through administration of the Edinburgh Handedness Inventory; Oldfield, 1971) with normal or corrected-to-normal vision were tested and reimbursed for their participation. Of these, three participants were excluded from the analyses, either because they were incapable of actively coping with the experimenters’ instructions (one participant) or because the total number of responses for one of the two conditions was too low (>20% of total trials for that condition; two participants). These exclusions resulted in a group of 23 participants (14 F, mean age 26.2±4.7 years). Written informed consent was obtained from each participant in accordance with the 2013 Declaration of Helsinki. The Ethics Committee of the University of Verona approved the experimental protocol (protocol number 07/2024).

### Stimuli

Stimuli consisted of a circular array of six letters pseudorandomly chosen amongst 19 of the 21 letters of the Italian alphabet (we did not include letters “O” and “N” to avoid confusion with “Q” and “M”, respectively), with half of them located to the right and half to the left of a fixation cross positioned at the center of the screen (at an eccentricity of 1.7° of visual angle). The chosen font for the letters was OpenDyslexic (https://opendyslexic.org/) in bold, a typeface designed to reduce confusion between similar letters. The letters were silver-colored (luminance: 24.71 cd/m^2^; x = 0.3167; y = 0.3399), each extending 1.8°, and were presented on a black 17-inch LCD monitor (luminance: 0.13 cd/m^2^; x = 0.2896; y = 0.2944). Letters were prevented from appearing in two consecutive trials to avoid any unintended repetition effect. Two different auditory cues at 2000 Hz and 500 Hz were played through a Bluetooth speaker and used in a counterbalanced manner to indicate to the participant which side of the stimulus to report.

### Experimental procedure

Participants sat in a dark room, 57 cm from the monitor. They placed their head on a chin rest to maintain their eyes aligned with the screen center. Participants were also instructed to keep their gaze on the fixation cross throughout the experiment.

The experiment started with six demonstrative, slowed-down trials to show the participants how trials were structured and to help them become accustomed to the two auditory cues. Then, a practice block identical to the experimental blocks (except that EEG was not recorded) was administered to the participants to help them familiarize with the task. After that, the actual experiment started. It consisted of ten blocks, each comprising 60 trials, interleaved with short pauses to allow participants to rest.

Each trial started with the experimenter pressing a key on the keyboard. After a 500 ms fixation interval, the stimulus appeared around the fixation cross and remained visible for 100 ms. Immediately following stimulus presentation, one of the two auditory tones was played for 500 ms, signaling the participant which side of the stimulus to report. Participants then had to report the target letters to an experimenter sitting behind them, who registered the response via a backlit keyboard. This entailed that the trial duration and inter-trial interval were not fixed, as they depended on the time necessary for the participant to answer and for the experimenter to register the response. Once the response was registered, the experimenter started a new trial by pressing a keyboard key (Fig. 1A).

**Figure 1.**
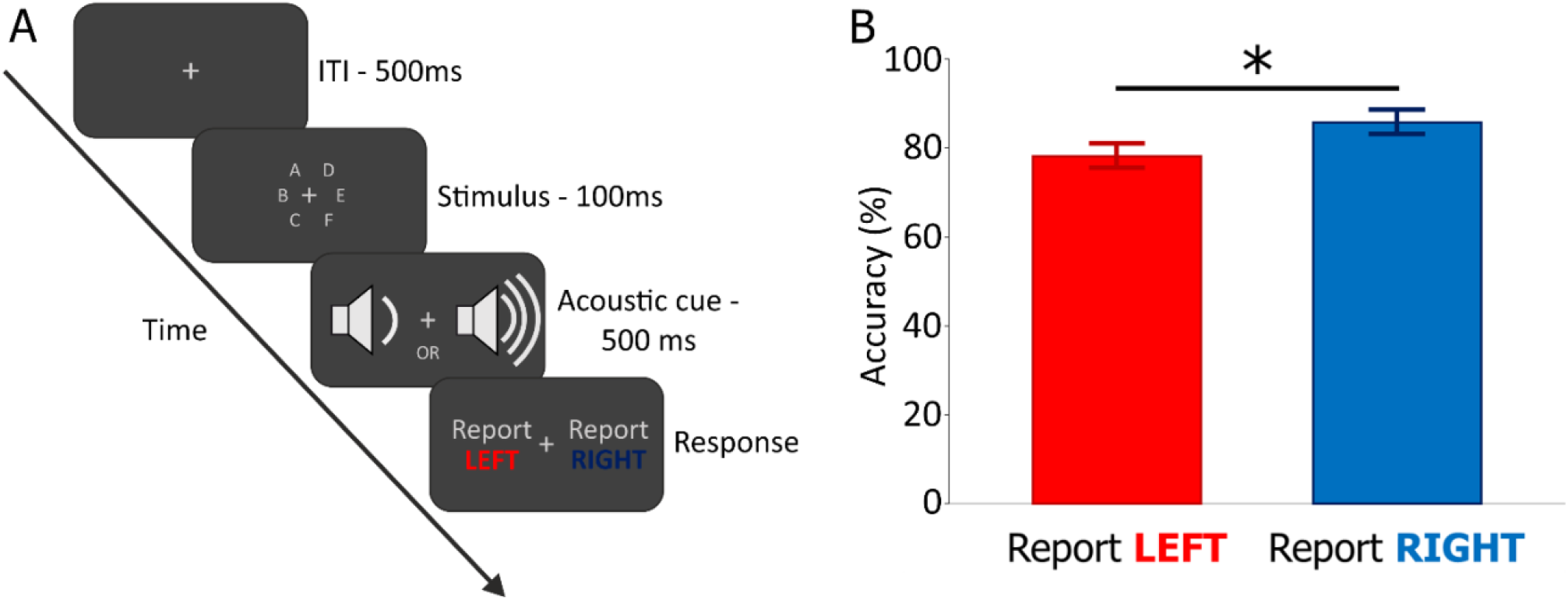
A: Outline of the experimental trial. B: Accuracy results for the two reporting conditions, “Report Left” (red) and “Report Right” (blue).

### EEG recording and preprocessing

To record the EEG signal, the software used was BrainVision Recorder, in combination with BrainAmp amplifiers (Brain Products GmbH, Germany) and a Fast’n East Cap (EasyCap GmbH, Germany) equipped with 59 passive Ag/AgCl electrodes placed according to the 10-10 International System. Six additional electrodes were used as online reference (RM), ground (AFz), and to control for vertical and horizontal eye movements. Electrode impedance was kept below 5 KΩ. The signal was digitally recorded at a sampling rate of 1000 Hz, with a 250 Hz low-pass filter and a 10-second time constant as the low cut-off in AC mode.

The EEG signal was preprocessed offline with EEGLAB (version 2024.0) (Delorme & Makeig, 2004) running in Matlab 2022b (Mathworks, USA). The signal was first downsampled at 500 Hz and high-pass filtered at 0.5 Hz (zero-phase, 3300^th^-order non-causal FIR band-pass filter). Then, the continuous data were epoched from 500 ms before to 2000 ms after the stimulus onset. Only trials in which participants reported three correct letters out of three were retained. On these, we performed a baseline correction using the time interval from -500 ms to 0 ms. Then, a manual trial rejection eliminated those trials contaminated by huge noise levels. Independent component analysis (ICA) was then used to remove brain-unrelated components from the data (e.g., ocular artifacts, muscle activity, electrode noise) (Delorme et al., 2007). On average, we removed 5.1±3.5 components. We then applied a notch filter (49-51 Hz, zero-phase, 1650^th^-order non-causal FIR band-stop filter) and a low-pass filter (70 Hz, zero-phase, 96^th^-order non-causal FIR band-pass filter). We average-referenced the data and performed another baseline correction (-500 to 0 ms time interval). As a final step, we performed a manual trial rejection to identify any poor-quality trials that might still contaminate the signal, and we epoched the data from -300 ms to 1500 ms.

### Statistical analyses

The software JASP was used to analyze behavioral data statistically (JASP Team, 2020). We calculated accuracy as the number of correctly reported letters per trial. Accuracy was determined both for each side to report and on average.

To analyze EEG data, we used the STUDY function of EEGLAB to perform a series of parametric, point-by-point, paired sample t-tests on the whole epoch and channels, with *Side to report* (Left|Right) as the grouping variable and a significance threshold of 0.05. False Discovery Rate (FDR) was used to address the multiple comparison problem (Groppe et al., 2011). To perform ERP analyses, we only considered trials in which participants reported 3/3 correct letters.

To shed light on the relationship between electrophysiological activity and performance, we run a backward linear regression (removal threshold: p > 0.05) with the components’ amplitudes as predictors of participants’ accuracy. We also run a correlation between the TIF component (see *ERP results* in the Results section) and the number of intrusions – i.e., letters from the uncued side and mistakenly reported. Regarding the components’ amplitudes, because the exact peak timing was often slightly different between the two reporting conditions, we chose an intermediate timing between the two conditions’ peaks, compatible with our sampling rate. When an exactly intermediate time point was not available, we chose the immediately subsequent time point instead – e.g., the first component peaked at 90 ms in one condition and at 92 ms in the other, so we chose 92 ms as the latency, since 91 ms was not available given our sampling rate. Once the latency was determined, we calculated the average amplitude within ±30 ms of the peak for each component, except for the last TIF component, for which we used a ±50 ms time window. The electrodes used to calculate amplitude were selected based on topography, with no differences between the two reporting conditions except for the TIF component, for which ERP analysis revealed a lateralized pattern of activity. In this case, homologous electrodes were chosen for the two reporting conditions.

## Results

### Behavioral results

The mean accuracy for the “Report Left” condition was 78.2% ± 10% (minimum: 58.4%; maximum: 93.1%). The mean accuracy for the “Report Right” condition was 85.8% ± 7.4% (minimum: 72.8%; maximum: 95.4%). A paired samples t-test showed a significant difference between the two conditions [t(22) = -4.026; p < 0.001; Cohen’s *d* = -0.840] (Fig. 1B). On average, participants’ accuracy was 82% ± 7.5% (minimum: 70.2%; maximum: 93.2%).

### ERP results

On average, we performed ERP analyses on 152.1±50.1 trials for the “Report Left” condition and on 196.6±43.4 trials for the “Report Right” condition. The ERP elicited by the two reporting conditions showed a P1 component (90 ms for “Report Left”, 92 ms for “Report Right”), followed by an N1 (158 ms for “Report Left”, 160 ms for “Report Right”) and a P2 (242 ms for “Report Left”, 244 ms for “Report Right”). Our stimuli failed to elicit a clear N2 component, while we found a positive deflection in the P3 time window (336 ms for “Report Left”, 340 ms for “Report Right). The next component peaked over occipital and bilateral temporal electrodes (726 ms for “Report Left”, 734 ms for “Report Right”), followed by a large posterior deflection ipsilaterally lateralized with respect to the side to report (1050 ms for “Report Left”, 1070 ms for “Report Right”). Throughout this work, we refer to these components as “Visual Code Reactivation” (VCR) and “Task-dependent Information Filtering” (TIF), respectively (see the Discussion for further information).

As shown in Figure 2, statistical differences between the two reporting conditions were observed only in a time window from ∼850 ms to ∼1100 ms, i.e., around the TIF peak. More specifically, “Report Left” elicited higher activations over left occipital, parietal, and centro-parietal electrodes, while the effect of “Report Right” was more widespread on the scalp, including electrodes from parieto-occipital to frontal areas.

**Figure 2.**
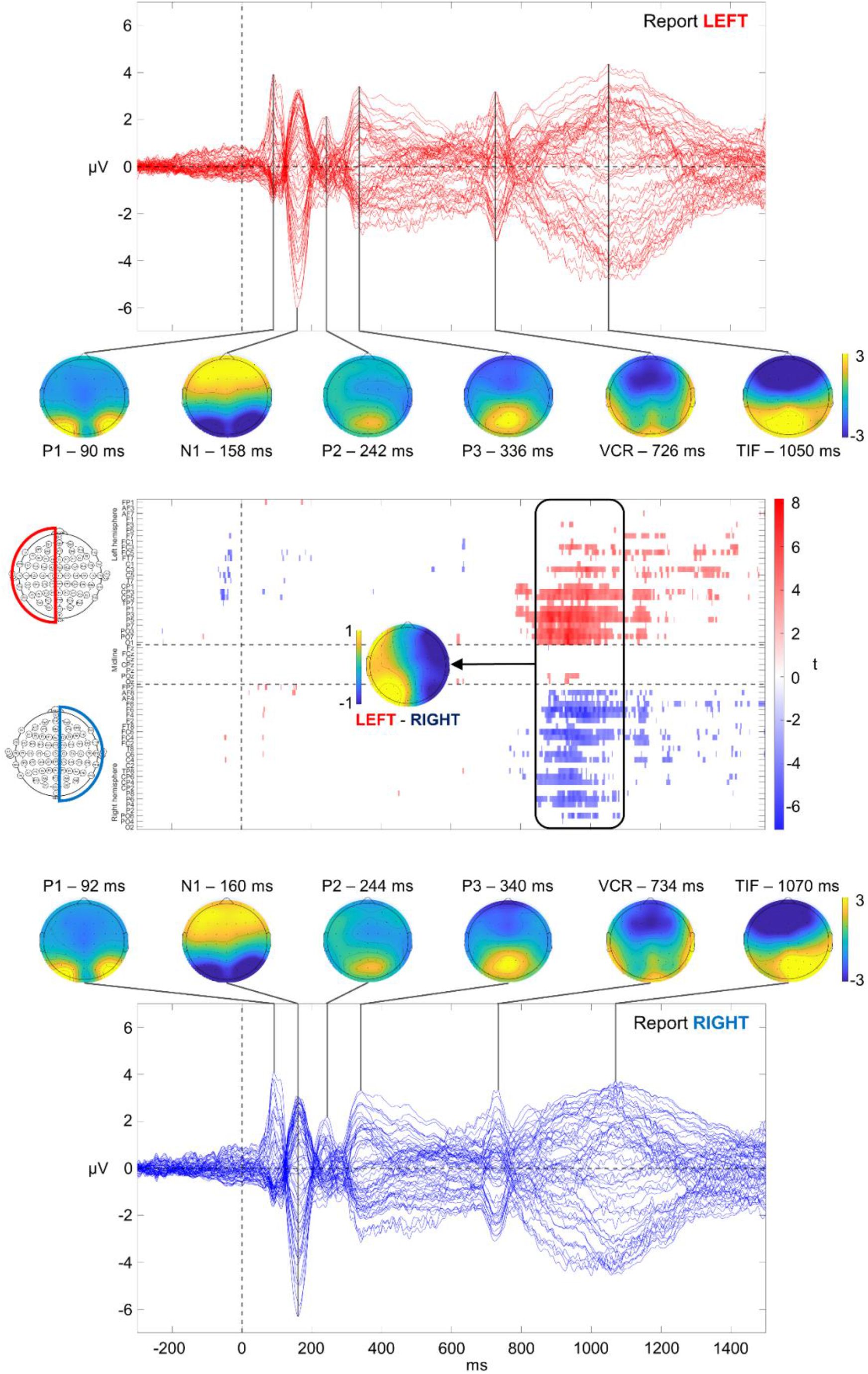
Butterfly plots, topographies, and raster plot for the two reporting conditions. The upper and lower sections contain the butterfly plots for the “Report Left” (red) and “Report Right” (blue) conditions, respectively. In addition, the topographies represent scalp distributions of the components’ peaks for the two conditions. In the middle, a raster plot shows, for each time point and each electrode, the significant differences (expressed as t values) between “Report Left” and “Report Right”. The black rectangle highlights the time window 850-1100, where statistical differences were more concentrated. The topography on its side represents the difference between the averaged “Report Left” and “Report Right” signals within this time window.

### Regression and correlation results

The backward elimination procedure resulted in three iterations before the final model was obtained (see Supplementary Material). The final regression model was statistically significant [F(1,4) = 11.968; p < 0.001], explaining ∼73% of the variance in accuracy (R^2^ = 0.727). Of the initial six predictors, only four of them significantly predicted accuracy in the final model: P1 (β = 0.387; t = 3.130; p = 0.006), P3 (β = -0.342; t = -2.356; p = 0.030), VCR (β = -0.313; t = -2.278; p = 0.035), and TIF (β = -0.439; t = -3.324; p = 0.004) (Fig. 3A-3D).

For the TIF, we found a positive correlation with intrusions (Pearson’s r = 0.421, p = 0.046) (Fig. 4).

**Figure 3.**
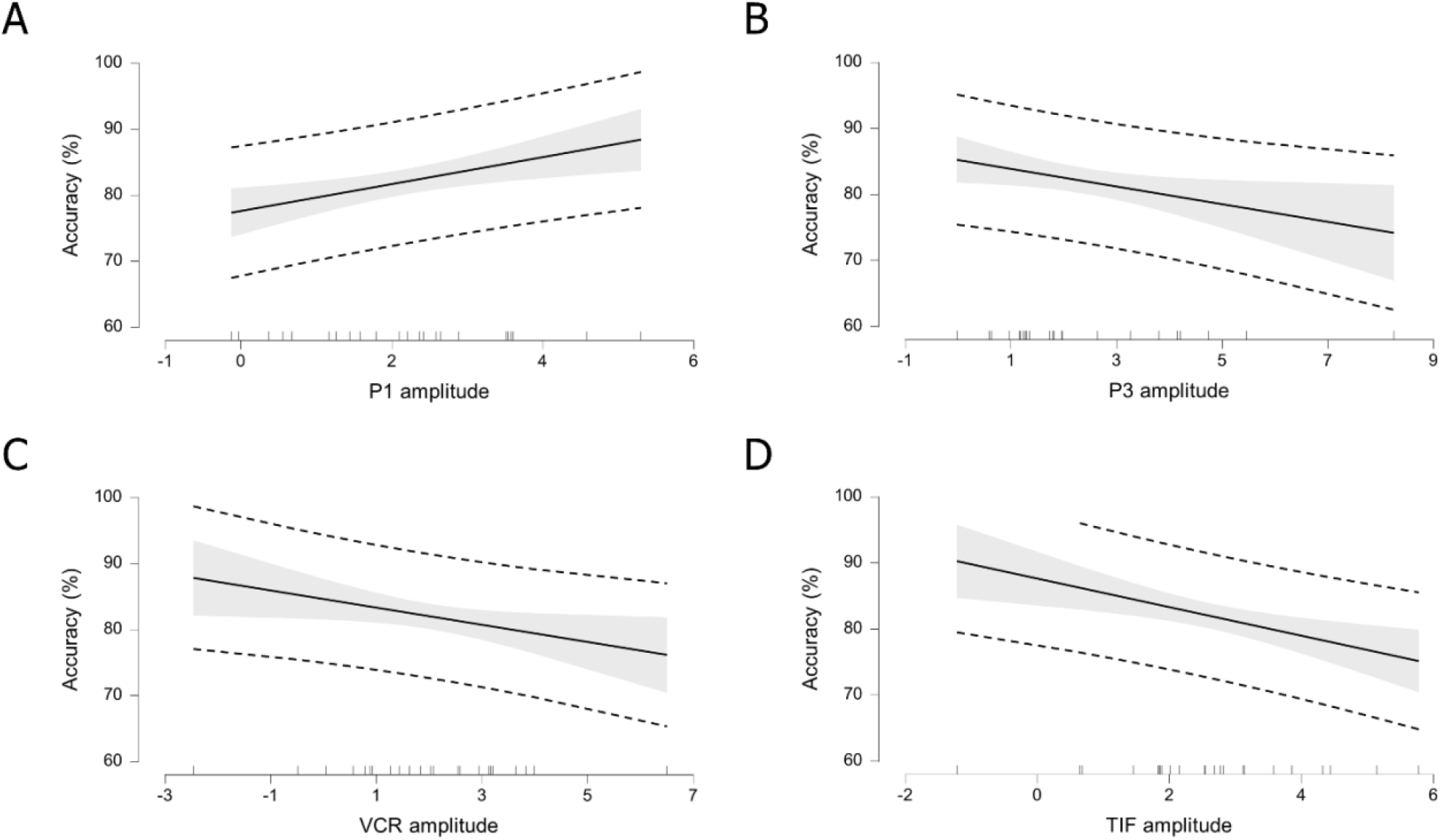
A-D: Marginal effects plots for the four significant predictors (P1, P3, VCR, TIF) on accuracy estimated from our multiple linear regression model. The solid black lines represent the impact of a one-unit change in each variable’s amplitude on accuracy, while keeping all other amplitudes constant. Shaded areas represent 95% confidence intervals, while dashed lines represent 95% prediction intervals.

**Figure 4.**
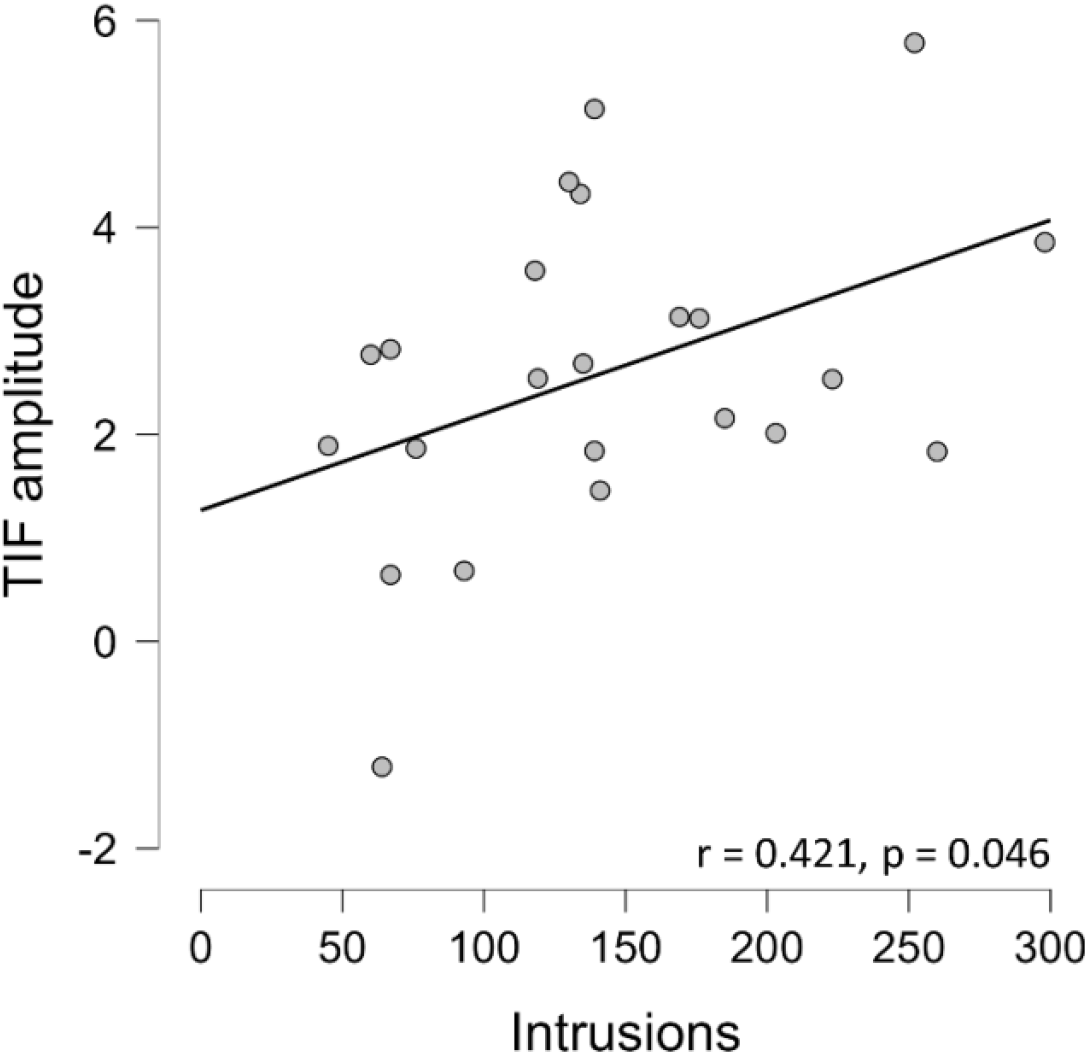
Scatter plot showing the positive correlation between the number of intrusions and TIF amplitude across participants. Each dot represents a participant. The solid line indicates the least-squares regression fit.

## Discussion

In this study, participants performed a partial report task while EEG was recorded. They were shown a letter array and had to report either the left or right half following a subsequent acoustic cue. This experiment was designed to disentangle the electrophysiological activations related to stimulus perception from those linked with post-perceptual processes. To do so, we took advantage of the properties of iconic memory, the early sensory store characterized by high capacity and very brief storage duration (Sperling, 1960). Accordingly, all letters should be present in iconic memory, independently of the side to report; however, as soon as the cue signals the relevant part to be reported, the electrophysiological activity related to subsequent post-perceptual processing should differentiate between the two reporting conditions, allowing for the isolation of the neural activity correlating with response creation.

Overall, our participants’ accuracy matched that of the original study (Sperling, 1960). It is worth noting that a recent replication study of Sperling’s (Bradley-Garcia et al., 2023) failed to reproduce these high accuracy levels, with the authors pointing to their participants’ relative lack of training. Considering our results, however, this seems unlikely: our participants achieved high accuracy levels with just 60 practice trials, roughly matching the number of practice trials in Bradley-Garcia’s and colleagues’ study.

When we looked at accuracy levels separately for the two reporting conditions, we found a significant difference in favor of the “Report Right” condition, which could be determined (in a non-mutually exclusive way) either by the superiority of the right visual field/left hemisphere in discriminating fine temporal events compared to the left visual field/right hemisphere (Nicholls, 1994; Nicholls & Atkinson, 1993; Nicholls & Cooper, 1991; Okubo & Nicholls, 2005, 2008) or by the left hemisphere superiority in processing language material (Brysbaert, 2004; Cohen et al., 2000; Dehaene et al., 2015; Ducrot & Grainger, 2007; Kermani et al., 2018; Levy et al., 2010; Rima et al., 2020).

To better understand the temporal unfolding of iconic memory, we examined the ERPs elicited by our partial report paradigm. This task could elicit a series of well-defined EEG components, with similar latencies in the two reporting conditions, reflecting successive stages of visual processing, from the initial reception of sensory information to the production of a stimulus-related response. The early sensory components (P1/N1) constitute the first broad elaboration of the visual stimulus (Pratt, 2011). The first elicited component was a P1, symmetrically located over parieto-occipital electrodes and likely representing the scalp projection of the occipital activations elicited by the two stimulus halves. It was followed by an N1 over most occipital and parietal electrodes, up to the centroparietal region. These early components may constitute the electrophysiological correlates of iconic memory: in vivo recordings in macaque monkeys have linked iconic memory to either the decaying post-stimulus response of V1 neurons (Teeuwen et al., 2021) or to the sustained activity of neurons in temporal areas (Keysers et al., 2005). These two hypotheses are not necessarily mutually exclusive, as they could capture two different moments of a highly recursive process (Ruff et al., 2007). Indeed, it is well known that visual processing relies on a series of feedforward, horizontal, and feedback loops (Lamme & Roelfsema, 2000). Whatever the case, it is noteworthy that both these candidate mechanisms occur within the first few hundred milliseconds, matching the latencies of our P1 and N1 components. The following P2 was characterized by a relatively small peak and a reduced extension, involving only the three most central parieto-occipital electrodes. This component could serve as a passageway from higher stages of visual processing (Omoto et al., 2010; Portella et al., 2012) to encoding for cognitive operations (Cepeda-Freyre et al., 2020; Finnigan et al., 2011; Lefebvre et al., 2005). Our stimuli did not elicit a visible N2, but we found a P3, consisting of a bilateral deflection over parieto-occipital, parietal, and centro-parietal electrodes. This component is classically associated with information processing from the sensory to the cognitive realm (Polich, 2011), including encoding of sensory-processed information (Busch & Herrmann, 2003). Here, the P3 component could represent the encoding of information from iconic memory into a more stable memory storage (Sligte et al., 2008), where it is retained until the acoustic cue is fully processed, a decision is made, and an output response is given. After the P3, we found a temporo-occipital positive deflection comprising medial occipital and lateralized temporal electrodes. This component, with its topography consistent with reactivation of higher visual areas (e.g., V4), may represent a re-encoding of the visuospatial traces associated with the encoded stimulus (Sligte et al., 2009). For this reason, we named it “Visual Code Reactivation” (VCR).

The following positive deflection extended from occipital to central electrodes. Importantly, this was the first component to exhibit a different topography for the two reporting conditions: the positive peak was lateralized ipsilaterally to the side of report - i.e., on left posterior electrodes for the “Report Left” condition, and on right posterior electrodes for the “Report Right” condition. This was further confirmed by our ERP analysis, showing that the two reporting conditions differed mainly at these electrodes in the time window from 850 to 1100 ms. This could represent a hemisphere-specific filtering mechanism preventing the irrelevant half of the reactivated stimulus from being recollected and reported. Because of this, we refer to this component as “Task-dependent Information Filtering” (TIF). Given its lateralized topography and speculative role, the TIF component presents conceptual similarities with the so-called “Distractor Positivity” (P_D_) (Gaspar et al., 2016; Gaspelin et al., 2023), typically investigated in visual search and working memory tasks. Likewise, TIF could reflect the process of inhibiting the irrelevant half-stimulus, allowing only the cued one to be correctly reported (Gaspelin & Luck, 2019).

This interpretation is based on the classic view of iconic memory. According to it, participants initially stored the entire stimulus, so that the neural activity did not differ between reporting conditions until they chose which side to report. Once this happened, participants had to filter out the irrelevant side, which determined the electrophysiological differences between the two conditions in the form of a TIF component.

To further characterize the elicited components, we ran a linear regression with the components’ amplitudes as predictors of participants’ accuracy. We found that four components were associated with performance: P1, P3, VCR, and TIF. P1 showed a positive association with accuracy, possibly stemming from its sensory nature: stimuli perceived more clearly led to stronger encoding and higher overall accuracy (Perri et al., 2014; Rutman et al., 2010). Conversely, P3, VCR, and TIF showed a negative association with accuracy: these relationships, according to the neural efficiency hypothesis (Haier et al., 1988, 1992) could reflect varying proficiency across the processing steps required by a partial report task, with more skilled individuals using fewer – and more focused – neural resources to cope with task demands (Li & Smith, 2021; Neubauer & Fink, 2009). This pattern has been confirmed not only by neuroimaging studies (Neubauer et al., 2002) but also by electrophysiological research in vision (Sanchez-Lopez et al., 2014) and memory (Herzmann & Curran, 2011).

We also found that intrusions positively correlated with TIF amplitude. This finding further reinforces the interpretation of this component as a filtering mechanism: lower amplitudes indicate that only relevant information is being further elaborated, while higher ones suggest that some unrequested letters made their way through, resulting in worse performance.

In recent years, Sperling’s original work on partial report and iconic memory has served as the cornerstone of the so-called overflow argument in consciousness research (Block, 2011; Stazicker, 2018). This states that perceptual systems provide rich contents, “overflowing” post-perceptual systems, characterized by smaller and sparser capacities, that access these perceptions to perform cognitive operations. This hypothesis draws on the distinction between two different concepts of consciousness (Block, 1990, 2005). The first, phenomenal consciousness, is the minimal neural basis for the content of a certain experience, the “what it is like” to have a certain experience (Nagel, 1974). The second, access consciousness, represents the conscious information as it becomes available for cognitive processing to the different brain systems, as if it were broadcast from a global workspace to the various subsystems (Dehaene & Changeux, 2011; Mashour et al., 2020). In the context of our experiment, all the stimulus letters enter phenomenal consciousness through iconic memory and are maintained in a pre-access modality until the cue is presented. After that, this phenomenal information must be accessed to retrieve the relevant letters: ERP differences between the two conditions might represent the first manifestation of access consciousness, with irrelevant letters filtered out while relevant ones undergo further processing. The first part of our elicited electrophysiological response, therefore, could be a neural correlate for the pre-cognitive, perceptually rich phenomenal consciousness, while later activations, differentiating between reporting conditions, are compatible with the idea of a selective, more restricted access for subsequent cognitive elaboration. Incidentally, the fact that relevant letters are maintained for a duration exceeding that of iconic memory but before being consciously accessed by working memory hints at the existence of an intermediate visual memory stage (Block, 2011; Sligte et al., 2008, 2009, 2010, 2011).

In conclusion, by exploiting the properties of iconic memory in a partial report paradigm, we were able to describe the entire processing flow of the visual stimulus, from the initial snapshot created by iconic memory to the selection of information for output. Critically, we provided evidence for a dissociation between early sensory encoding and the later formation of consciously reportable visual content, finding differences between reporting conditions only in this latter step and suggesting that, up to that point, the stimulus is entirely retained in the brain. Only when the side to report is determined is irrelevant information filtered out.

Future investigations should focus on further characterizing the neural correlates of iconic memory: for example, via source reconstruction or oscillatory analysis, or by adding a “full report” condition to further disentangle the information availability during early sensory processing.

## Supporting information

Suppelementary materials

## Supplementary data

Supplementary data is available as a separate file.

## Data availability

Data will be made available on reasonable request.

