## Supplementary material for "From Encoding To Conscious Report: Electrophysiological Signatures Of Iconic Memory Revealed By A Partial Report Task": Suppelementary materials

**Supplementary materials**

*Model Summary*

| | | | | | Durbin-Watson | | |
|---|---|---|---|---|---|---|---|
| Model | R | $R^2$ | Adjusted $R^2$ | RMSE | Autocorrelation | Statistic | p |
| $M_0$ | 0.858 | 0.735 | 0.636 | 4.563 | -0.075 | 2.112 | .856 |
| $M_1$ | 0.858 | 0.735 | 0.658 | 4.427 | -0.074 | 2.112 | .862 |
| $M_2$ | 0.852 | 0.727 | 0.666 | 4.372 | -0.031 | 2.020 | .980 |

Supplementary Table 1. Summary of the three iterations of the linear regression model. $M_0$ is the first iteration of the model, with all six components as predictors (P1, N1, P2, P3, VCR, TIF); $M_1$ is the second iteration of the model, with predictors (P1, N1, P3, VCR, TIF); $M_2$ is the third iteration of the model, with four predictors (P1, P3, VCR, TIF).

*ANOVA*

| Model | | Sum of Squares | df | Mean Square | F | p |
|---|---|---|---|---|---|---|
| $M_0$ | Regression | 925.765 | 6 | 154.294 | 7.411 | < .001 |
| | Residual | 333.094 | 16 | 20.818 | | |
| | Total | 1258.860 | 22 | | | |
| $M_1$ | Regression | 925.712 | 5 | 185.142 | 9.448 | < .001 |
| | Residual | 333.148 | 17 | 19.597 | | |
| | Total | 1258.860 | 22 | | | |
| $M_2$ | Regression | 914.861 | 4 | 228.715 | 11.968 | < .001 |
| | Residual | 343.998 | 18 | 19.111 | | |
| | Total | 1258.860 | 22 | | | |

Supplementary Table 2. Summary of the three ANOVAs performed, one for each iteration of the model.

*Coefficients*

| Model | | Unstandardized | Standard Error | Standardized | t | p |
|---|---|---|---|---|---|---|
| $M_0$ | (Intercept) | 87.741 | 3.161 | | 27.754 | < .001 |
| | P1 | 2.148 | 0.714 | 0.408 | 3.008 | .008 |
| | N1 | -0.209 | 0.294 | -0.100 | -0.710 | .488 |
| | P2 | 0.035 | 0.699 | 0.011 | 0.051 | .960 |
| | P3 | -1.382 | 0.884 | -0.353 | -1.563 | .138 |
| | VCR | -1.252 | 0.603 | -0.301 | -2.078 | .054 |
| | TIF | -2.053 | 0.716 | -0.418 | -2.868 | .011 |
| $M_1$ | (Intercept) | 87.737 | 3.066 | | 28.613 | < .001 |
| | P1 | 2.156 | 0.678 | 0.410 | 3.179 | .005 |
| | N1 | -0.205 | 0.275 | -0.098 | -0.744 | .467 |
| | P3 | -1.349 | 0.576 | -0.344 | -2.342 | .032 |
| | VCR | -1.255 | 0.583 | -0.302 | -2.153 | .046 |
| | TIF | -2.064 | 0.667 | -0.420 | -3.094 | .007 |
| $M_2$ | (Intercept) | 89.023 | 2.501 | | 35.599 | < .001 |
| | P1 | 2.035 | 0.650 | 0.387 | 3.130 | .006 |
| | P3 | -1.339 | 0.569 | -0.342 | -2.356 | .030 |
| | VCR | -1.303 | 0.572 | -0.313 | -2.278 | .035 |
| | TIF | -2.153 | 0.648 | -0.439 | -3.324 | .004 |

Supplementary Table 3. Summary of the information related to intercept and coefficients of the model, including unstandardized values, standard error, standardized values, and corresponding t-values and p-values.
